# Cryo-EM reveals alternative modes of dimerization driving activation of IKK

**DOI:** 10.64898/2026.06.29.735262

**Authors:** Tapan Biswas, Shandy Shahabi, Xiang-Yang Zhong, Myung Soo Ko, Tom Huxford, Gourisankar Ghosh

## Abstract

The inhibitor of κB kinase (IKK) complex integrates diverse cellular inflammatory responses, and induces transcription factor NF-κB. The molecular mechanism by which IKK becomes catalytically active in response to signaling remains unclear despite structural knowledge of the individual IKK1/α, IKK2/β, and NEMO/IKKγ protein components within its hetero-oligomeric assembly. Cryo-EM of the IKK2/β homodimer bound to an associating NEMO/IKKγ protein fragment, reveals multiple conformers. Mutual exclusivity of dimeric conformers, canonical versus alternate, is reflected in and dependent upon order-to-disorder transition of the canonical 6-helical bundle dimerization interface. Correlation of this unusual structural plasticity of IKK2/β with its biochemical and cellular activities suggests mechanistic possibilities for how association with its partner scaffold protein NEMO/IKKγ and polyubiquitin chains might dictate catalytic activation of IKK through distinct IKK2/β conformers.

**Significance:** The inhibitor of κB kinase (IKK) complex is central to inflammatory signaling via the NF-κB family transcription factors. Its activation mechanism has remained unclear. Cryo-EM analysis reveals that the constituent kinase IKK2/β adopts structurally distinct, mutually exclusive dimeric conformations controlled by an ordered-to-disordered transition at its canonical dimerization interface. Stabilization of select IKK2/β conformers by the scaffold protein NEMO in association with poly-ubiquitin chains is regulated through modular architecture and structural plasticity of distinctive kinase-associated domains, present only in kinases of this family. This unique regulatory mechanism governing catalytic activation of IKK2/β provides a conceptual framework for targeting dysregulated NF-κB signaling in human diseases.

---

IKK2 (also known as IKKβ), a kinase domain-containing subunit of the IKK complex, is one of the central regulators of inflammation and innate immunity. Together with its paralog IKK1 (IKKα), IKK2 emerged during the evolution of multicellular organisms alongside its essential accessory subunit NEMO (NF-κB essential modulator; also known as IKKγ) (1-6). The N-terminal kinase domain (KD) of IKK1 and -2 displays all the conserved signature features of canonical Ser/Thr protein kinases (**Fig. 1A**)(7-9). This pair of kinases, along with TBK1 and IKKε, belong to a unique structural subgroup in which the KD is followed in sequence and structure by a ubiquitin-like domain (ULD), and a scaffold dimerization domain (SDD) composed of three long α-helices forming a triple helical bundle (10-15). The contribution of the ULD and SDD to IKK activity has thus far remained enigmatic, except that a region of the SDD that is distal to the KD has been observed to mediate homodimerization in experimentally determined structures of IKK1 and IKK2(16). Additionally, a small α-helical NEMO-binding domain (NBD) is appended at the extreme C-terminal end of IKK1 and IKK2 through a long, disordered linker (16).

**Fig. 1.**
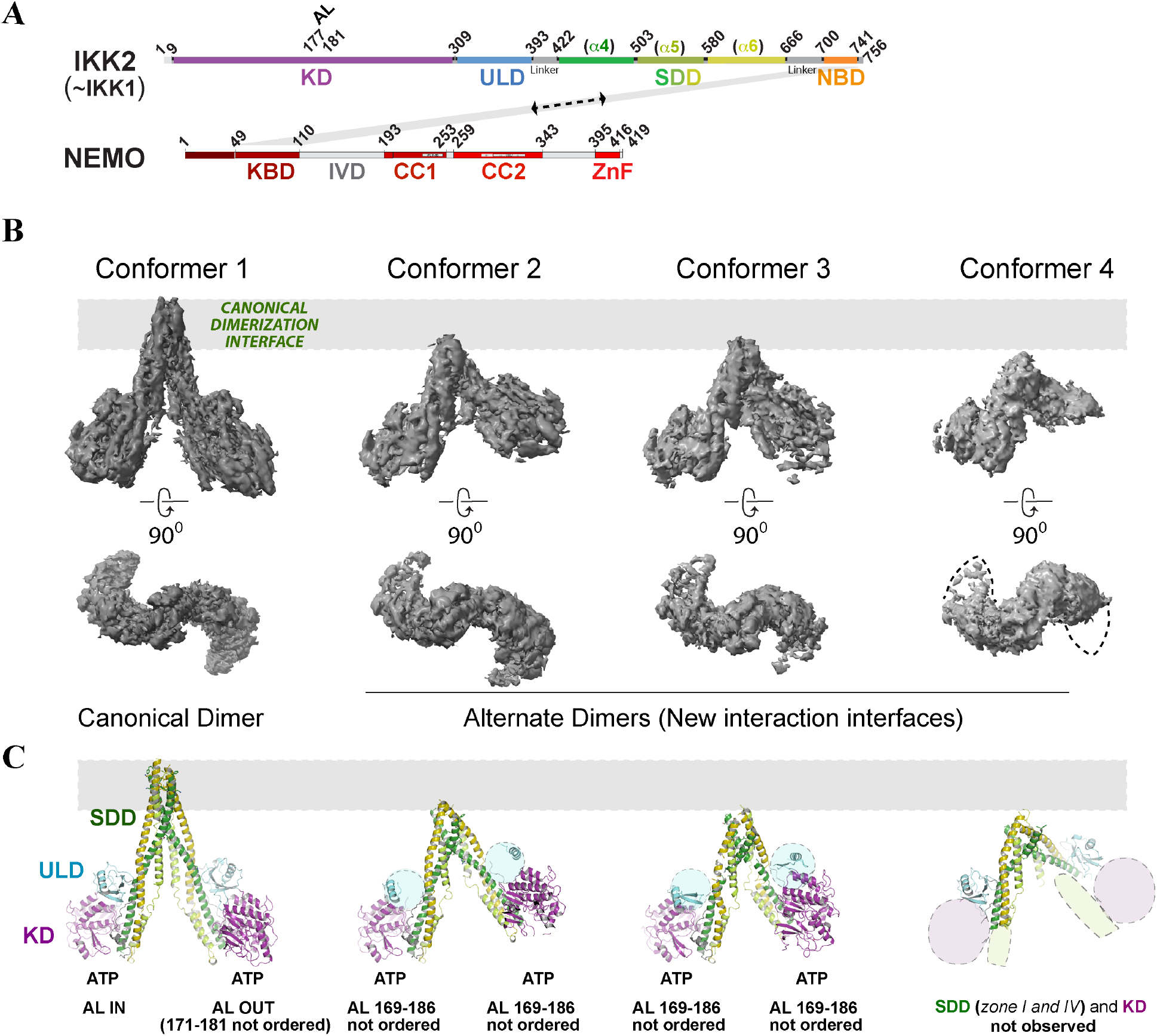
Structural plasticity of human IKK2. **A**, A schematic of the organization of IKK1/2 and NEMO domains. Residues at the domain boundaries and relevant elements involved in activation are indicated. **B**, cryo-EM maps (orthogonal views) of IKK2 homodimer in presence of NEMO^1-110^ (contour level 0.25) indicating four distinct conformers with alternate dimerization modes. **C**, Models of IKK2 homodimers built using the electron potential maps, showing a canonical and three alternate conformers. Pronounced asymmetry is observed in the alternate dimers as a result of the flexible SDD, displaced ULD, and dislodged KD. The domains are color coded as indicated in (a) and missing regions are highlighted with shaded ellipses. The presence of ATP in the catalytic pocket and activation loop features are noted below. Gray bars highlight the SDD region missing in alternate conformers.

Despite their high degree of global structural similarity, IKK1 and IKK2 are quite different in many aspects of their activation and functionality, and they selectively trigger distinct transcriptional programs through different sets of NF-κB family transcription factors (17-19). IKK1 plays a dominant role in immune cell differentiation and development, while IKK2 is rapidly activated in response to diverse pro-inflammatory and pathogenic stimuli, triggering inflammatory and cellular survival responses. Catalytic activation of both IKK1 and IKK2 requires, and is defined by, phosphorylation at two serine residues (S176 and S180 of IKK1; S177 and S181 of IKK2) located within the activation loop (AL) of their respective KD (2, 8). One key factor underlying this stimulus-dependent and temporally controlled activation of IKK2 is the generation and interaction of linear and/or branched polyubiquitin chains (Ub-chain) with NEMO (20).

The X-ray crystal structures of the core KD-ULD-SDD domains of IKK1 and IKK2 serve as valuable models from which to untangle the structural basis for IKK catalytic activity and substrate specificity (10-13). In addition, crystallographic models of the IKK1 and IKK2 NBD complexed with the kinase binding domain (KBD) of NEMO (NEMO^1-110^) reveal the basis for tight association between the three main components of the IKK complex (16). Despite this hard won experimental structural data, mechanistic insights into the stimulus-dependent activation of the IKK complex are just beginning to be unraveled. In this study, we analyzed the structure of full-length IKK2 bound to NEMO^1-110^ by cryo-EM, which revealed an unexpected structural plasticity of IKK2 homodimerization, underlying its regulated activation.

## NEMO-bound IKK2 displays a heterogenous population of dimers dominated by the canonical conformer

Purified full-length IKK2, expressed in recombinant baculovirus-infected Sf9 insect cells, was complexed with purified NEMO^1-110^, expressed in *E. coli*, in the presence of Mg-ATP. The resultant complex was applied to plasma-cleaned Quantifoil Au grids for vitrification, and 19,626 frames were collected on a 300 kV TEM (Titan Krios; ThermoFisher) fitted with an energy filter and a Falcon IV detector (**Fig. S1**). Heterogenous classification of particles forming good 2D-class averages (**Fig. S2**) revealed multiple dimeric architectural forms of IKK2, with four major distinct 3D-conformational states (conformers 1-4; **Fig. 1B, Figs. S3, S4, S5, and Table S1**) – each comprising of ∼150K to ∼550K particles. However, the NBD:KBD complex tetra-helix region, previously characterized by X-ray crystallography, was not observed in any of resultant 3D reconstructions.

The predominant species (conformer 1) closely resembled the previously structurally characterized IKK2 homodimer, in which dimerization is mediated by a region of the SDD distal to the KD. The individual protomers of this IKK2 homodimer are largely symmetrical; however, although both contain ATP bound at their nucleotide binding sites, their respective AL regions adopt clearly distinct conformations (details in Fig. 4A). This suggests a functional nonequivalence of the protomers within the dimer that is independent of their interaction with ATP (**Fig. 1C**).

The other conformers (2, 3, and 4) deviate markedly from the canonical architecture of conformer 1, and, although homodimeric, conspicuously lack the canonical dimerization interface. Instead, these IKK2 homodimers are stabilized through different inter-protomer contacts of alternate dimerization interfaces, and are hereafter referred to as the “alternate” dimers. Conformers 2 and 3 share a similar overall architecture and a common alternative dimer interface. As with conformer 1, both protomers of conformers 2 and 3 bound ATP similarly. However, the AL segment 169-186 of both protomers is largely unresolved, and the density corresponding to the ULD domain is largely absent.

Conformer 4 displays pronounced conformational variability throughout the molecule. Although the SDD helices at the alternate dimer interface and the apposed ULDs are relatively well-resolved, the KDs and the associated proximal SDD regions are highly variable (diffuse density in the maps), and thus the presence or absence of ATP could not be ascertained. The absence of the KD-proximal region of SDD reflects the necessity of KD for its conformational stability. Although the dimeric interface of conformer 4 bears similarity to that of conformers 2 and 3, the interfacial surface area is reduced and the SDD helices are more frayed: clearly divergent from the interface of conformer 1.

## Alternate conformers reveal conditional instability of the canonical dimer interface

Analysis of the four IKK2 conformers observed via cryo-EM of IKK2:NEMO^1-110^ complexes reveals that the signature triple-helical SDD of IKK2 consists of smaller modular “zones,” each with independent interaction and unique structural/functional properties that appear to contribute toward regulation of IKK complex activity (**Fig. 2A,B**). Helix-α4^422-503^ (IKK2 residues 422-503) and helix-α5^504-580^ are interrupted at specific locations, whereas helix-α6^581-666^ remains uninterrupted. Helix-α5^504-580^ seems to be particularly flexible, as evidenced by its decreased resolution in maps (**Fig. 2C, Fig. S6**).

**Fig. 2.**
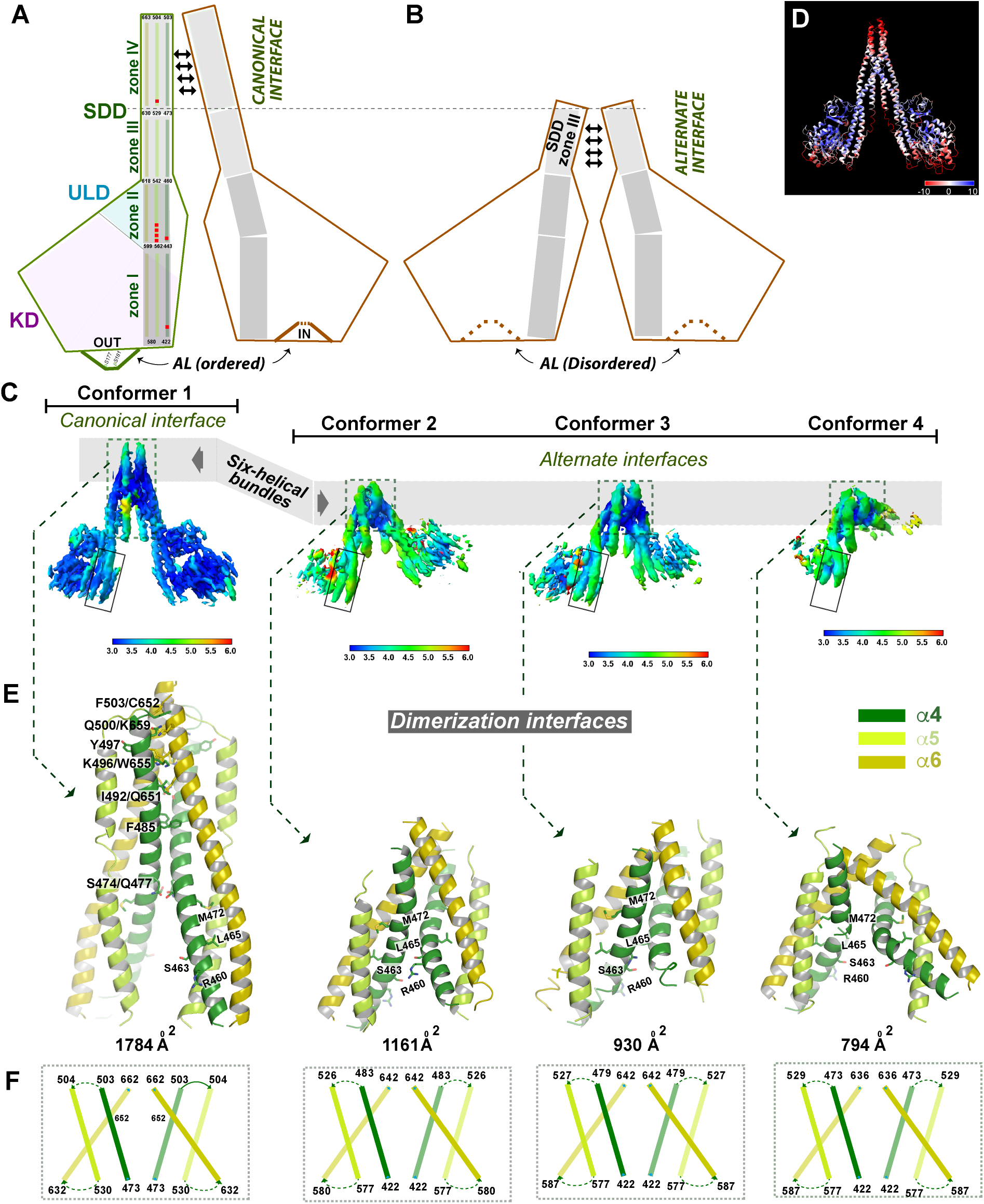
Dimer interfaces in canonical and alternate conformers of IKK2. **A**,**B** Schematics of canonical and alternate conformers highlighting the zonal subdivisions of SDD, and positioning/nature of AL loops. Red dots mark locations of glycine residues and flexible regions. **C**. Electron potential maps of canonical and alternate IKK2 homodimers (contour level of 0.6 to highlight helix path) surface colored by local resolution and aligned using SDD zone I of one protomer as a frame of reference (solid rectangle) highlights the alternate interfaces attainable upon dissolution of the canonical interface. **D**, Higher flexibility of helices at the canonical interface and peripheral zones displayed in B-factor map of the canonical model from refinement using REFMAC Servalcat with half-maps. **E**, Details of the helices forming canonical and alternate dimerization interfaces (key residues are highlighted); buried surface area is indicated below. **F**, A schematic of the helices at the dimerization interface with end residues marked.

The region of the SDD proximal to the KD (zones I and II), corresponding to residues 422– 460 of helix-α4^422-503^, grazes the KD and ULD, knitting them together, while its most distal segment (zone IV) contributes to formation of the canonical dimer interface. Zone IV helical parts, α4^473-503^, α5^504-529^, and α6^630-663^ from both of the protomers form a composite 6-helical bundle, which appears to be dynamic as reflected by higher temperature factors within this region (**Fig. 2D**). The alternate dimers (conformers 2–4) deviate from the canonical (conformer 1), differing primarily through structural rearrangements of the helices around SDD zones II and III (IKK2 residues 443-473). Within the density maps of each conformer, subclasses with global variations are evident, which could not be reliably segregated into distinct 3D classes. Thus, these conformers belong to a structural continuum rather than discrete and well-separated states. Despite this conformational heterogeneity, the relative positioning of six SDD helices from two protomers, as well as the positions of the KDs and ULDs relative to the SDDs, are quite clear. The central role of the loose packing among the long SDD helices and their internal interruptions becomes apparent, as these features allow switching among mutually exclusive alternative and canonical dimer interfaces that are spatially close and orientationally constrained. Key determinants to this remodeling are the pronounced flexibility of residues 550–559, together with the helix-breaking glycine residues at position 525 in helix-α5^504–580^ and at positions 431 and 450 in helix-α4^422-503^ (**Fig. 2A**). The 550–559 region acts as a hinge to allow both shift and rotation of the SDD zone II (residues 543–561 and juxtaposed helical parts) relative to the SDD zone I (562–580 and juxtaposed parts).

The dimer interfaces of conformers 2 and 3, which are only permissible as a consequence of shift and rotation between individual zones by SDD helices α4, α5, and α6, are similar in overall location of the interacting regions of the SDD helices and the interaction surface (**Fig. 2E, F**). These two conformers, which both lack a structurally ordered SDD zone IV region, represent an end of a structural continuum opposite to that of the conformer 1. The SDD zone III residues of helix-α4 (∼456-473) that are solvent exposed in conformer 1, are juxtaposed against α4 and α6 of the neighboring protomer to form the alternate dimer interface. The surface area buried at the alternate dimer interfaces, although significantly reduced, is nonetheless substantial compared to that of the canonical dimer interface, suggesting stability (**Fig. 2E**). It is noteworthy that the combined population of particles with alternate dimer interfaces is comparable to those with the canonical dimer interface (750k vs 550k during early stages of heterogeneous classification; **Fig. S3**).

Overall, Zone I of the IKK2 SDD contacts the KD and provides mutual stability; zone II interacts with the ULD to establish a communication channel between this region of the SDD and the KD; zone III provides an alternative homodimer interface in the absence of canonical dimerization, which keeps protomers within close proximity, albeit in an inactive state; and zone IV forms the canonical dimer and directs the interplay of different domains in IKK2 for its timely phosphorylation of specific substrates. These findings reveal a surprising plasticity of SDD and suggest how zonal subdivision of the SDD allows remodeling of IKK2 between alternate and canonical dimerization states. We postulate that regulation of this equilibrium among IKK2 conformers by NEMO in complex with Ub-chains is the crux of IKK2 activation.

## The SDD regulates structural states and activation of IKK2

The zone IV of the IKK2 SDD, which forms the canonical dimer interface, is disordered in a recent X-ray structure of an unpaired IKK2 protomer, indicating the requirement of dimerization for SDD zone IV to remain structured(21). A rearrangement of SDD helices in individual protomers of the three alternate dimeric conformers is necessary for disordered-to-ordered transition of the canonical dimerization interface (**Fig. 3A**). A consequence of this rearrangement is a slight displacement of the ULD with respect to the KD, and the consequent movement of the KD with its associated SDD zone I such that the KD achieves its stable canonical (active) arrangement (details presented in Fig. 4A).

**Fig. 3.**
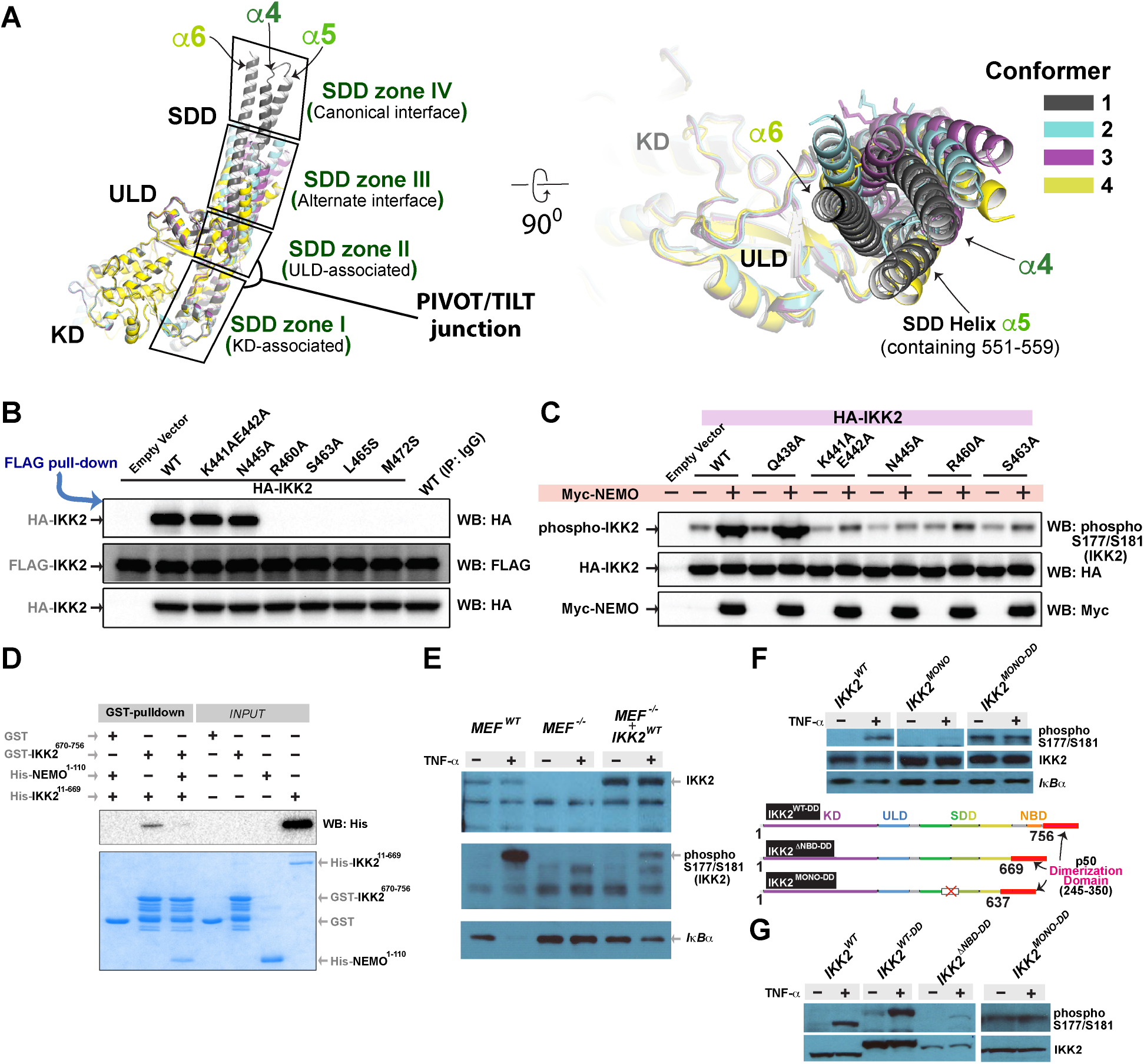
SDD Structural plasticity regulates dimerization modes and activation of IKK2. **A**. Superposed cartoons of four conformers of IKK2-dimer (orthogonal views) displaying positional variability of SDD independent modules (zone I to zone IV; black rectangles). The zone IV forming canonical interface is absent in alternate conformers. The zone III provides the interface for dimerization in alternate conformers. Zone II pivots with respect to the KD-apposed zone I and regulate positional constraints on ULD and KD. **B**. Expression plasmids of wild-type IKK2 (and its mutants) fused to either Flag or HA were co-transfected in HEK293T cells. Immunoprecipitation with anti-Flag antibody followed by WB with anti-HA implicate a role for specific residues inIKK2 dimerization through the alternate interface. **C**. Contributing role of residues at and near the alternative interface in AL phosphorylation of IKK2. Expression constructs of HA-tagged IKK2 (wild-type and mutant) were co-transfected with Myc-tagged NEMO in HEK293T cells. Induction of AL phosphorylation was assessed 24-h post transfection upon stimulation with TNF-α for 30 min. Residues mediating second contact between NEMO and IKK2 were also defective in activation, as reported previously. **D**. GST-pulldown assay showing interaction between IKK2^ΔNBD^ (residues 670-756) and IKK2^1-669^ and the detrimental effect of NEMO^KBD^ (residues 1-110). **E**. Reconstitution of IKK2^WT^ in ikk2^-/-^ MEF cells restores TNF-α-induced signaling. **F**. IKK2 deletion and fusion constructs showing requirement of alternative dimer interface in AL phosphorylation. **G**. AL phosphorylation of IKK2^ΔNBD^ and IKK2^MONO^ fused to a heterologous dimerization domain indicate that both canonical and alternative dimerization interfaces are required for inducible activation of IKK2 (Note: Inducibility IKK2^ΔNBD^ could not be tested due to poor expression).

**Fig. 4.**
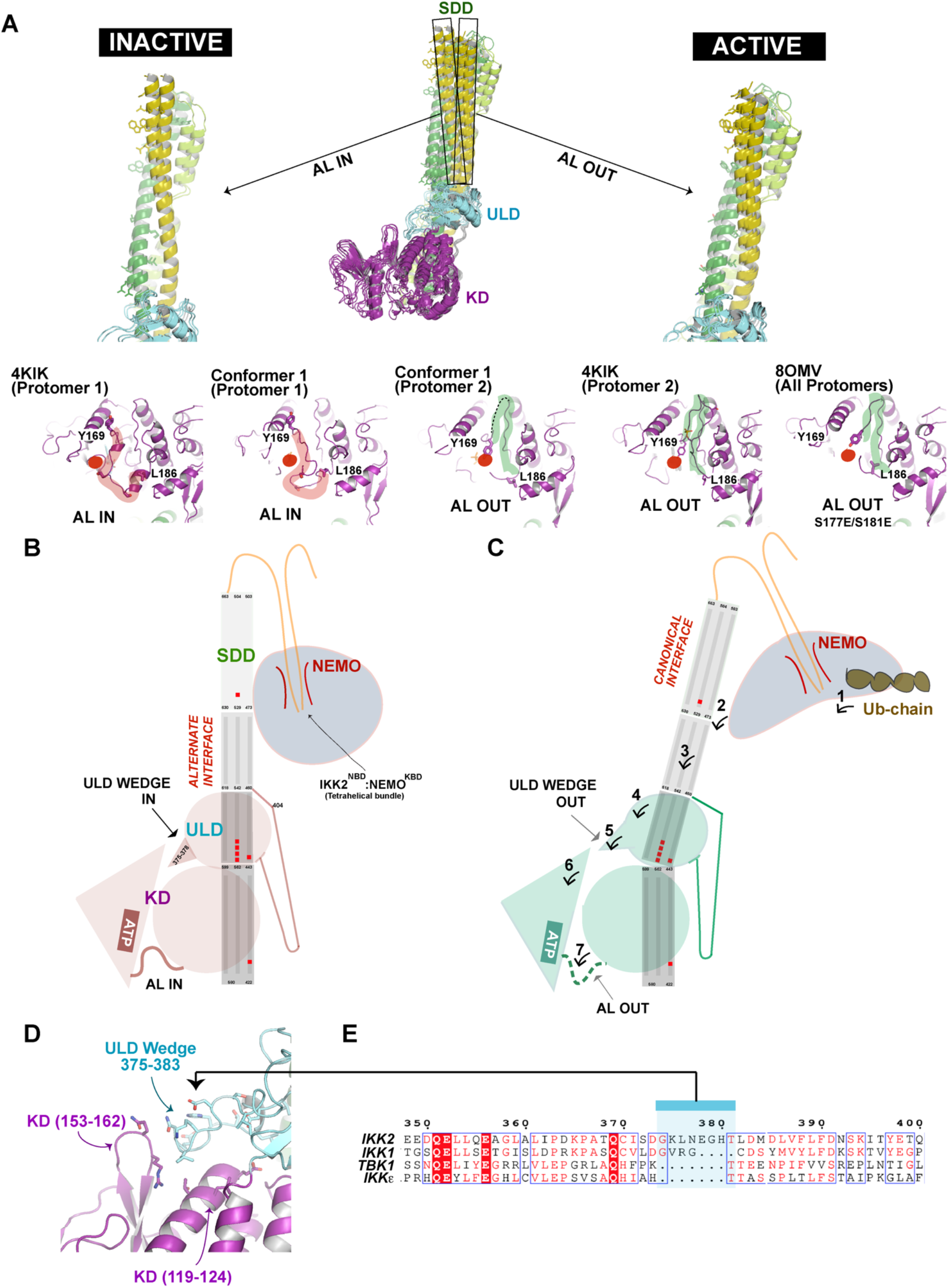
Dimerization-controlled activation of IKK2 occurs through an allosteric cascade of signals propagated via NBD to SDD to ULD to KD. **A**. Correlation between position of SDD helices and configuration of the AL. The tilt of the SDD helices with respect to the KD-ULD scaffold is distinct in the two protomers of the dimer, and determines the IN and OUT conformations of the activation loop. In all protomers of the 8OMV model (IKK2^S177ES181E^), the tilt matches with the active protomer of the canonical conformer (conformer 1) or the model 4KIK. Red circles indicate the ATP binding pocket of the protomers. The general path of the AL IN and OUT conformations are highlighted with red and green ribbons, respectively. **B**. Schematic of IKK2 in its inactive state with domain positions indicated. The four SDD zones that mediate the canonical interface, alternate interface, ULD-associated interface, and the KD-associated interface are highlighted. The ULD with its KD-interacting Wedge and the AL positions (IN and OUT) are also indicated. **C**. Activation of IKK2: Interaction of Ub-chain (1), reorganizes NEMO and regulates conformation of the IKK2:NEMO complex (2), promoting formation of dimers with canonical interface through SDD zone IV (3). The SDD reorganization repositions the ULD (4), and controls its interaction with KD through ULD Wedge residues 375-383 (5), reorganizing KD N- and C-lobe interface (6), and in turn ejects the AL (7) to accommodate substrate phosphorylation. Points of flexibility in SDD helices are marked with red dots. **D**. SDD transmits its reorganization effect to KD through an ULD Wedge. **E**. The ULD Wedge is unique to IKK2 and not present in other IKK subgroup members IKK1, TBK1, and IKKε

Prior to its structural characterization by X-ray crystallography, a purported leucine zipper domain spanning L458 to F486 of IKK2 was predicted to mediate subunit dimerization (2), and mutation of M472 was shown to impair IKK2 activation (2, 22). M472 resides within zone III of SDD helical bundle. Using immunoprecipitation, we observed association between co-expressed Flag-IKK2^WT^ and HA-IKK2^WT^; however, the HA-IKK2^M472S^ mutant failed to associate with Flag-IKK2^WT^ (**Fig. 3B**), consistent with previous reports. Additionally, we tested roles of residues located at or near the alternative dimer interface using mutants IKK2^R460A^, IKK2^S463A^, and IKK2^L465S^. L465 is a core-forming residue, like M472, within the helical bundle, R460 projects towards the ULD, and S463 lies directly at the subunit dimer interface. These variants also displayed defective dimerization similar to IKK2^M472S^ (**Fig. 3B**). These results hint at direct interactive roles of R460 and S463 and possibly an indirect role for L465 and M472 on structural stability of the helical bundle. We also examined HA-IKK2^K441A/E442A^ and HA-IKK2^N445A^ mutants as controls since these residues are not at the alternative dimer interface but play an important role in IKK2 activation by facilitating Ub-chain binding-dependent AL phosphorylation [17]. As anticipated, these mutants are not observed to be defective in protomer association (**Fig. 3B**).

We next assessed if defects in alternative dimerization could impair IKK2 activation. AL phosphorylation of IKK2 was measured by Western blot using an antibody specific for IKK2-phosphoS177/S181 in extracts of HEK293T cells overexpressing specific HA-IKK2 constructs in the presence and absence of NEMO (**Fig. 3C**). Both HA-IKK2^R460A^ and HA-IKK2^S463A^ mutants displayed defective AL phosphorylation regardless of overexpressed NEMO, further supporting the role of the alternative dimer interface in IKK2 activation. As we reported previously, HA-IKK2^K441A/E442A^ and HA-IKK2^N445A^ mutants also showed defective AL phosphorylation because of their unique role in communication with NEMO when NEMO binds to linear Ub-chains. In contrast, the HA-IKK2^Q338A^ mutant, which targets a region of IKK2 that is neither involved in alternative dimerization nor in Ub-chain-dependent binding of NEMO to IKK2, exhibited no defect in IKK2 activation (23). Collectively, these observations strongly support a novel regulatory role of SDD zone III-mediated alternative homodimerization in IKK2 activation, and hint at a functional coupling of NEMO and/or Ub-chain engagement to dimerization modes.

The likely influence of the NBD:KBD tetra-helical bundle in selecting distinct dimeric interfaces, and its conspicuous absence in density maps prompted us to test if the C-terminal NBD region of IKK2 interacts intramolecularly with the main body of the enzyme and, if so, whether this interaction is NEMO-dependent. We indeed observed an interaction between IKK2^666-756^ and IKK2^11-666^ (**Fig. 3D**). Furthermore, this interaction was negated via competition upon engagement of NEMO-KBD to IKK2-NBD. This strongly suggests a NEMO binding-dependent reorganization of IKK2, possibly involving its independent SDD modules.

To further verify the premise that dimerization is critical to regulated activation of IKK2, we generated several of its deletion (IKK2^ΔNBD^, IKK2^MONO^) and fusion (IKK^WT-DD^, IKK2^ΔNBD-DD^, IKK2^MONO-DD^) constructs, shown schematically in **Fig. 3F** (24). The fusion constructs contain the dimerization domain (residues 245-358) of the NF-κB p50 subunit. IKK^-/-^ MEF cells were reconstituted with wild-type (WT) and mutant IKK2 constructs. The WT MEF cells expressing endogenous IKK2 and the IKK2^-/-^ MEF cells reconstituted with IKK2^WT^ displayed TNF-α-induced AL phosphorylation of native and reconstituted IKK2 and IκBα degradation, as expected (**Fig. 3E**). However, the mutants revealed distinct regulatory requirements. No AL phosphorylation of IKK2^MONO^ was observed when reconstituted in MEF-IKK2^-/-^ cells, and only a negligible amount appeared in response to treatment with TNF-α. However, the IKK2^MONO-DD^ exhibited constitutive activation (**Fig. 3F, G**). This suggests that keeping the IKK2^MONO^ protomers in proximity through fusion to a heterologous DD promotes dimeric arrangement and subsequent structural changes that are conducive to AL phosphorylation. IKK2^ΔNBD-DD^ demonstrated activation upon induction, the rationale for which is unclear.

## A mechanistic path to IKK2 activation

To investigate possibilities of how the novel IKK2 homodimer interface could facilitate activation of IKK2, we analyzed differences among individual protomers within the four IKK2 dimeric conformers in this cryo-EM study, and in previously determined crystallographic models (PDB ID 4KIK and 8OMV). The protomers were aligned using KD-associated SDD zone I as the frame of reference, since we observed major translational and rotational shifts of SDD zone II relative to zone I in the different conformers. This revealed a surprising correlation between the position of the AL and the orientations of helices that constitute SDD zone II (**Fig. 4A**). An outward tilt (relative to dimer axis) of SDD zone II appears to transmit its effect on the interacting ULD, and through the ULD “Wedge” (IKK2 residues 375-383) to the KD and eject the AL (**Fig. 4B,C**). Prior literature indicates two distinct primary conformations for the AL, referred to as the “IN” and “OUT” (inactive and active, respectively). The two critical serines of the IKK2 activation loop – when either phosphorylated or substituted with phospho-mimetic glutamate residues (IKK2^S177E-S181E^) – are unequivocally established to adopt the AL OUT conformation, which permits access of substrates to the catalytic pocket and is associated with a catalytically active protein kinase. The structural model of the IKK2 canonical homodimer (conformer 1) in our study revealed asymmetric disposition of its two activation loops. A previously determined homodimer model (PDB ID: 4KIK) exhibited a similar asymmetry in which the AL of one protomer adopted the IN conformation with AL serines unphosphorylated, while the other adopted the OUT conformation with AL serines phosphorylated [18]. In both protomers of conformers 2 and 3, density corresponding to AL residues 169-186 was absent, and pronounced rotational/translation shifts of SDD zone III biased towards its counterparts in the AL IN (inactive) protomer of the conformer I. Steric clashes of ULD and KD were also evident in reconstruction of these alternate conformers. Thus, these conformations are likely reflective of earlier states in the potential pathway leading to activation, where the IKK2 domains/subdomains are not stably organized yet, imposing a higher flexibility to the active site and substrate-interacting region. The adoption of the canonical dimer conformation appears to allow for AL phosphorylation, which enables the kinase for productive substrate (e.g., IκB) binding and catalytic activity (**Fig. 4C**) (21).

Our study provides compelling evidence that heretofore unrecognized SDD-enabled dynamic transitions between modes of IKK2 homodimerization allow the conformational changes within the IKK2 subunit that are necessary for AL phosphorylation and consequent catalytic activation. The IKK2-NBD and the NEMO-KBD tetra-helical bundle, despite not being visible, facilitate proximity of IKK2 protomers but allow toggling among alternative homodimerization modes. Full-length NEMO likely maintains IKK2 in an inactive state via an unidentified region downstream of the KBD. This inhibitory activity of NEMO is relieved upon non-covalent binding of Ub-chains to the ubiquitin binding motif of NEMO, allowing transition of IKK2 into an activation-competent state. A NEMO peptide spanning residues 374-389, that mediate a Ub-chain induced secondary interaction with IKK2, has been shown to play critical roles in this process(23). Overall, this study points to a unique modular architecture of the IKK2 SDD that allow switching of the alternate homodimer interfaces to canonical – in turn inducing positional changes of the ULD and promoting the AL OUT conformation that enables phosphorylation of S177/S181. Thus, the ULD appears to function as a transducer of dimerization-dependent IKK2 activation (**Fig. 4D**). A comparison of the IKK2 to IKK1, and also to the TBK1 and IKKε, reveal that the ULD Wedge that we observed docked into the KD is unique for IKK2 (**Fig. 4E**). The absence of this transducer in IKK1 correlates with its predominant NEMO-independent functionality. These observations also hint at the intriguing relevance of IKK existing primarily as a heterodimer (IKK1:IKK2) in cell, likely stemming from differences at residues I492L, K496R, and F503Y leading to a stronger canonical interface. Moreover, IKK1 will obviously not adopt conformational transitions as the IKK2 due to key differences in ULD-KD-‘SDD zones I and II’ module, highlighted by the ULD Wedge. IKK-related proteins TBK1, which associates with optineurin, and IKKε also lack this ULD Wedge, consistent with a quite different interprotomer organization within the dimer and an even different regulatory mode mediated by the STING peptide(25, 26). Exploration of these distinct aspects of these kinases could enable us to target them selectively.

## Acknowledgments

The authors like to thank the UCSD cryo-EM facility for help with data collection.

## Author contributions

Conceptualization: TB, GG Methodology: TB, GG

Investigation: TB, SS, XZ, MSK, GG Visualization: TB

Funding acquisition: TH, GG Project administration: GG Supervision: TB, TH, GG

Writing – original draft: TB, TH, GG Writing – review & editing: TB, TH, GG

## Data availability

All data in the study are available in the main text, the supplementary materials, or in the public repository. The atomic coordinates and sharpened cryo-EM and half maps of various conformations of IKK2 dimers (Extended Data Table 1) have been deposited in the PDB (https://www.rcsb.org/structure/) and EMDB. The accession codes are: Canonical (Conformer 1) PDB 12MM and EMD-76572, respectively; Alternative (Conformer 2) PDB 12MN and EMD-76573, respectively; Alternative (Conformer 3) PDB 12MO and EMD-76574, respectively; Alternative (Conformer 4) PDB 12MP and EMD-76575, respectively.

