## Supporting Data for "Cryo-EM reveals alternative modes of dimerization driving activation of IKK"

**Figures S1-6 and Table S1:**

**a.** **b.**

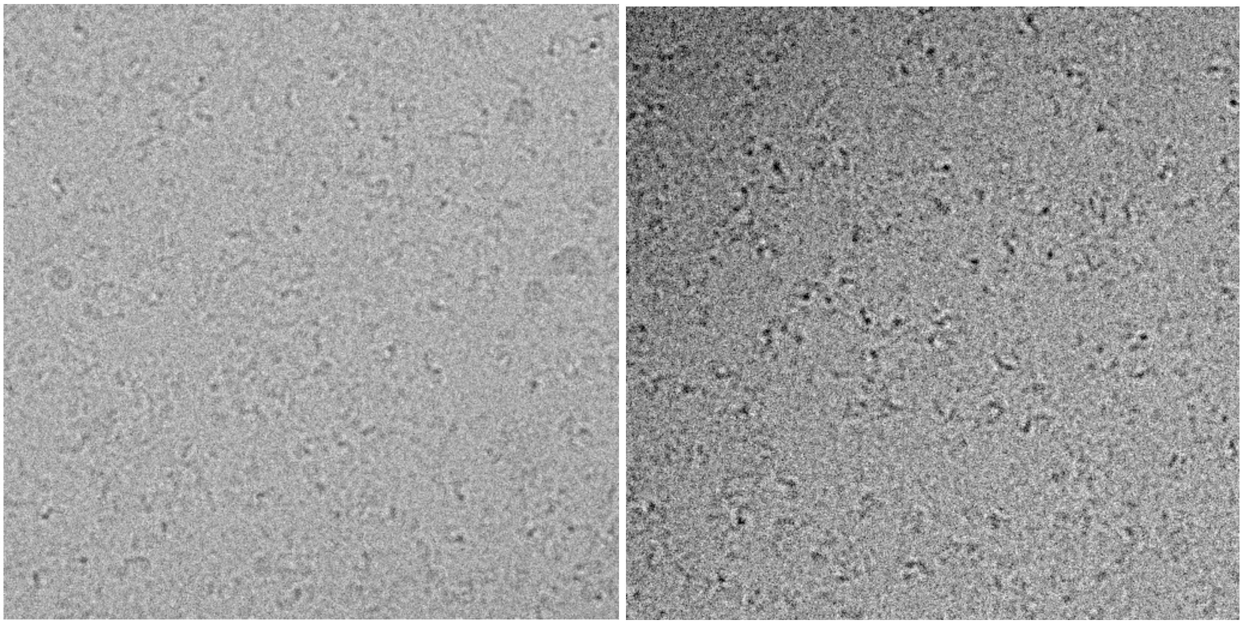

**Fig. S1.** Representative images of IKK2 and NEMO$^{1-110}$ complexes. **a.** at 0.8 μm defocus, **b.** 1.5 μm defocus.

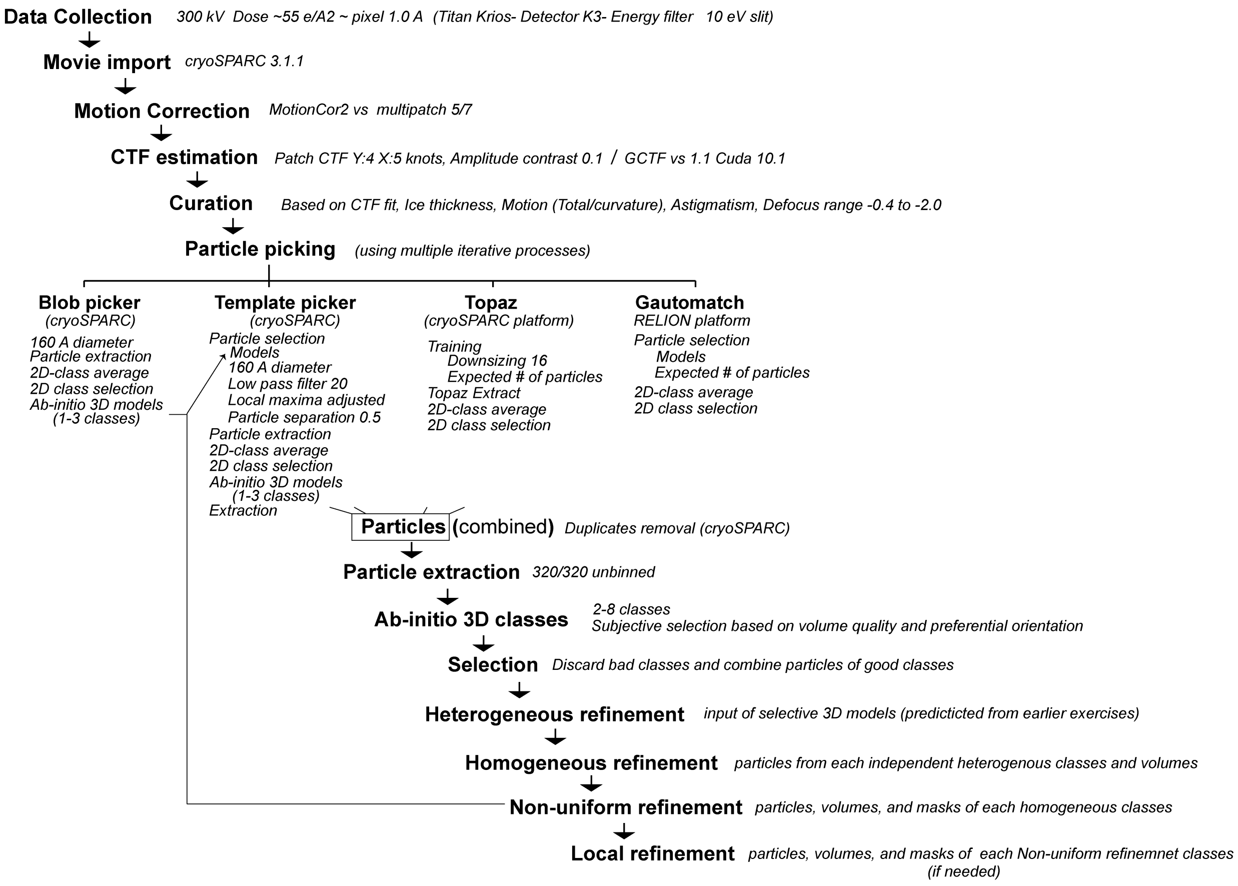

**Fig. S2.** Pipeline for cryo-EM data processing.

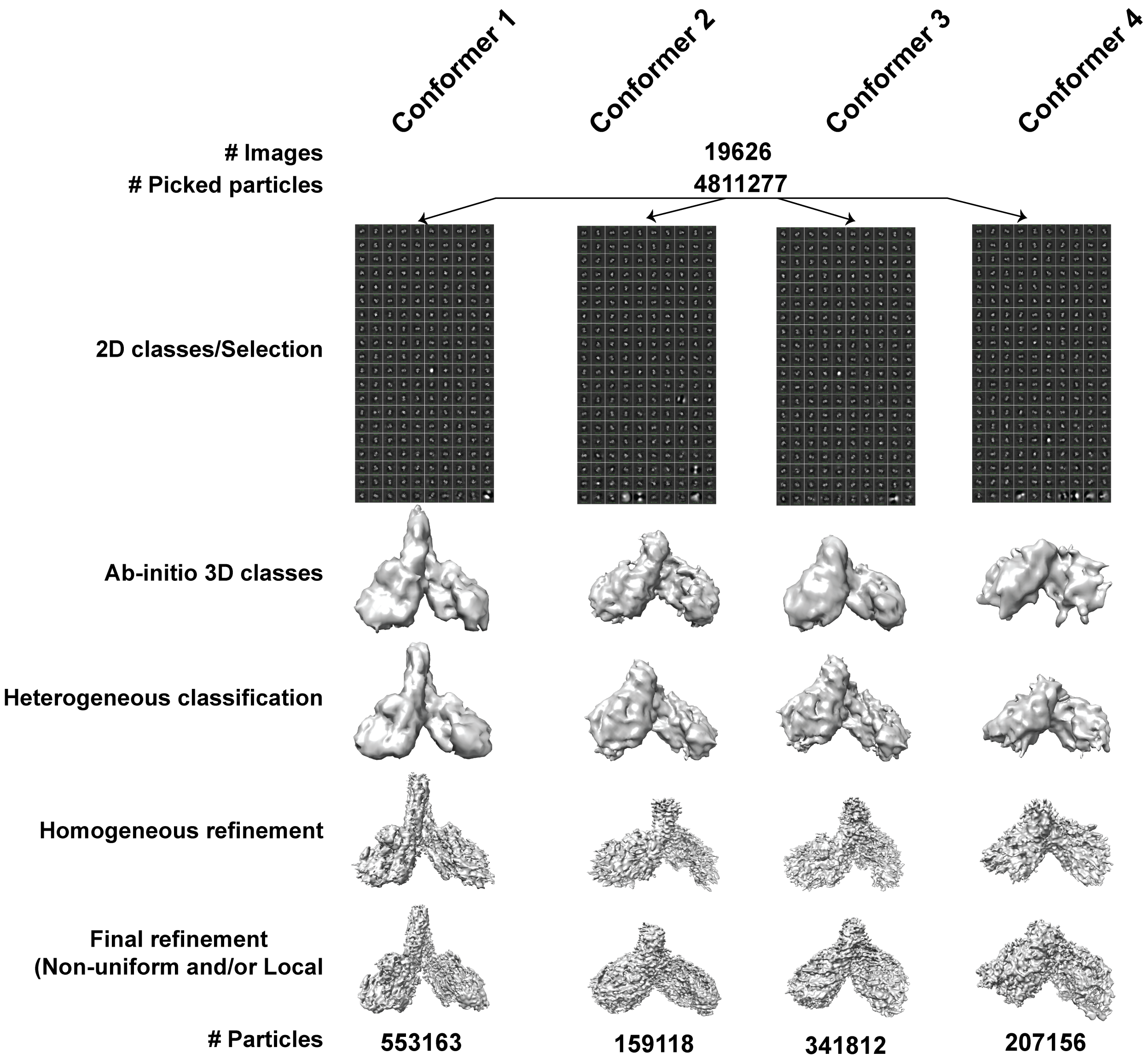

**Fig. S3.** Pipeline for map generation.

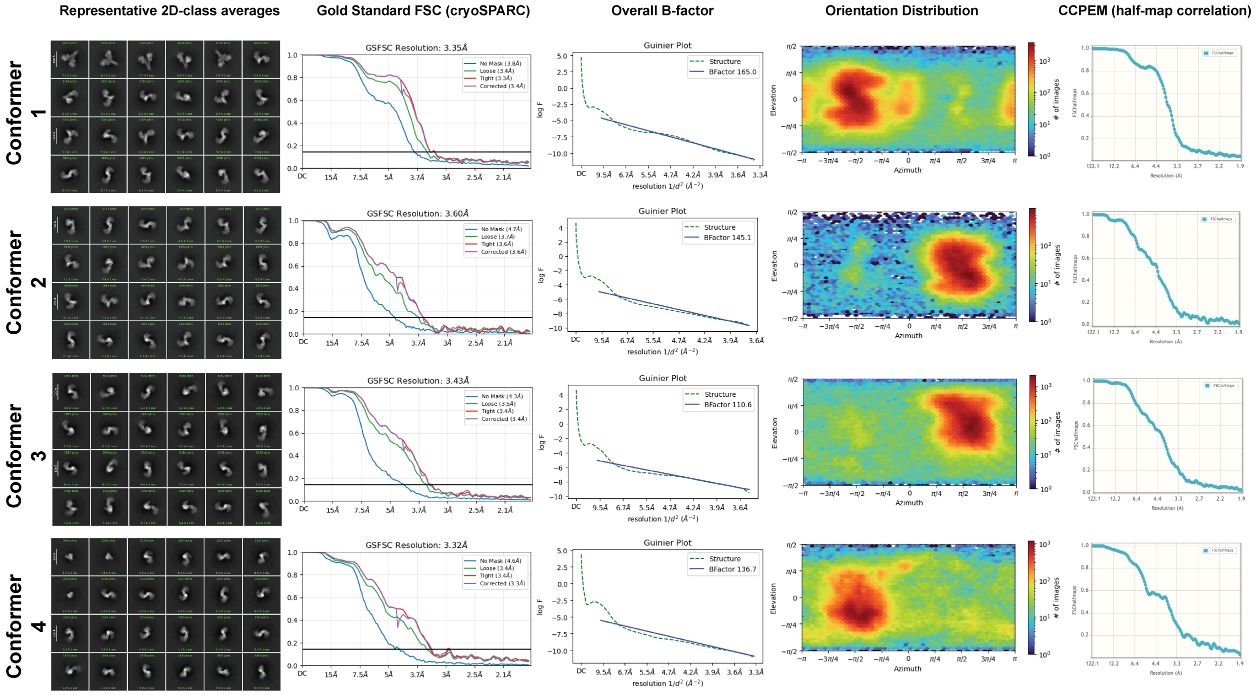

**Fig. S4.** Details of four unique conformers of IKK2 complex. Sets of horizontal panels indicate 2D class averages, cryoSPARC generated GSFSC curves, global B-factor of map, orientation distribution, and CCPEM derived half map correlation of four unique IKK2 models. The complexes are numbered as in Extended Data Table 1.

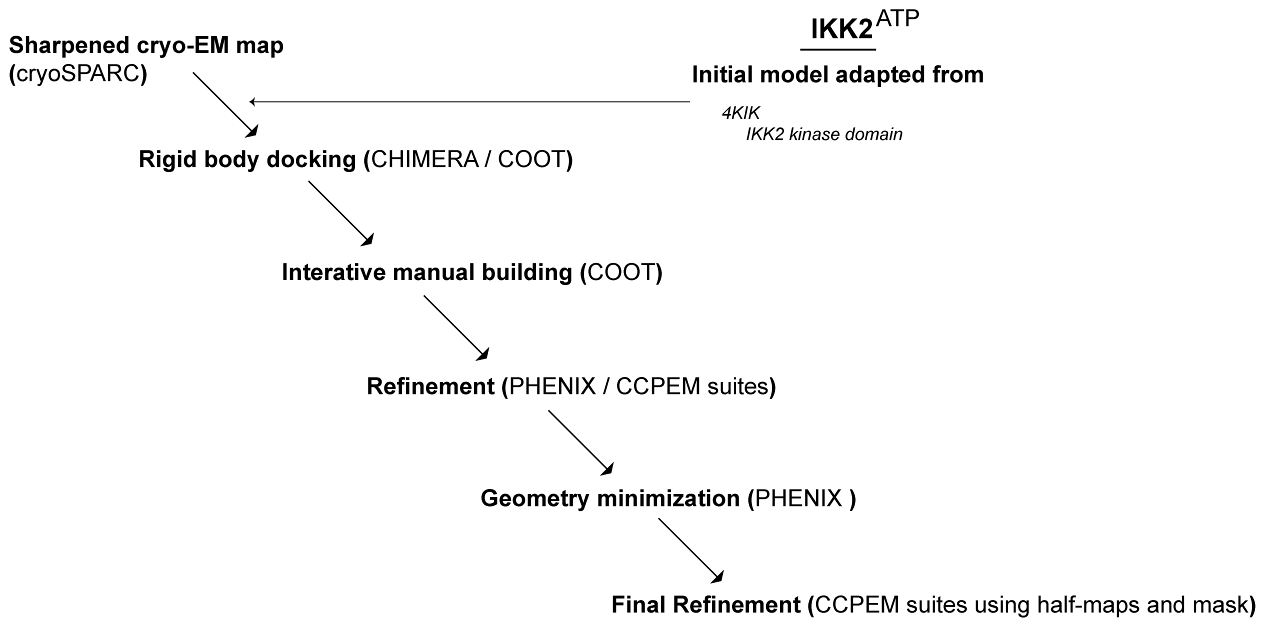

**Fig. S5.** A schematic of the atomic model building in the cryo-EM maps and refinement processes for IKK2 complexes.

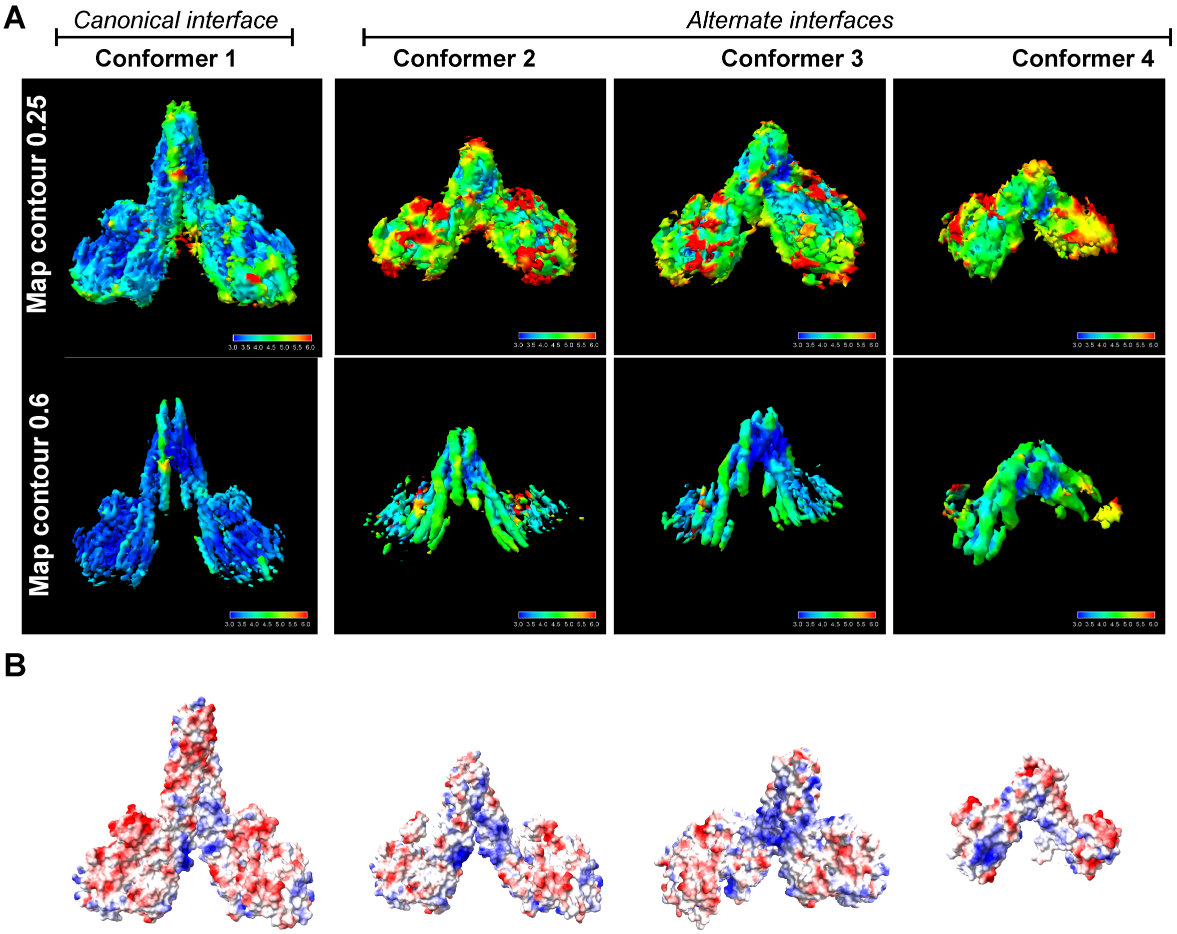

**Fig. S6. a.** Local resolution map at contour level 0.25 (map density snugly encompassing the model) and 0.6 (map density showing clarity of the helix backbones); **b.** surface charge properties of the four unique conformers of IKK2 complexes.

**Table S1.** Cryo-EM map and model coordinate statistics.

| Table 1. Cryo-EM data collection, refinement and validation statistics | | | | |
|---|---|---|---|---|
| IKK2-NEMO(1-110) Complexes | 1 | 2 | 3 | 4 |
| Conformation | **Conformer 1** | **Conformer 2** | **Conformer 3** | **Conformer 4** |
| Dimeric Interface | Canonical | Alternate | Alternate | Alternate |
| Incubation | ATP | ATP | ATP | ATP (not visible) |
| Map (EMDB code) | **EMD-76572** | **EMD-76573** | **EMD-76574** | **EMD-76575** |
| **Data Collection and Processing** | | | | |
| Magnification | 130,000 | 130,000 | 130,000 | 130,000 |
| Voltage (kV) | 300 | 300 | 300 | 300 |
| Electron exposure (e-/A2) | ~ 55 | ~ 55 | ~ 55 | ~ 55 |
| Defocus range (nm) | 400-2000 | 400-2000 | 400-2000 | 400-2000 |
| Map size (A) | 320/320/320 | 320/320/320 | 320/320/320 | 320/320/320 |
| Pixel size (A) | 0.935 | 0.935 | 0.935 | 0.935 |
| Total images used (#) | 19626 | 19626 | 19626 | 19626 |
| Final particle images (#) | 465567 | 127422 | 148084 | 109161 |
| Global map resolution (A) | 3.35 | 3.6 | 3.43 | 3.32 |
| FSC threshold | 0.143 | 0.143 | 0.143 | 0.143 |
| Global B factor (map) | 165 | 145.1 | 110.6 | 136.7 |
| **Model composition** | | | | |
| Model (PDB code) | **12MM** | **12MN** | **12MO** | **12MP** |
| Total atoms (non hydrogen) | 10574 | 8406 | 8524 | 3245 |
| Protein residues/atoms | 1301/10512 | 1027/8344 | 1043/8462 | 410/3245 |
| Ligand atoms (ATP) | 62 | 62 | 62 | none |
| **RMS deviation** | | | | |
| Bond lengths (RMSZ) | 0.08 | 0.11 | 0.08 | 0.07 |
| Bond angles (RMSZ) | 0.23 | 0.25 | 0.24 | 0.22 |
| **Validation** | | | | |
| Molprobity Clash score (including H atoms) | 3 | 4 | 4 | 3 |
| Clashes | 74/10574 | 69/8406 | 76/8524 | 20/3245 |
| Ramachandran plot (protein backbone) | | | | |
| Favored (%) | 93 | 94 | 92 | 96 |
| Allowed (%) | 6 | 5 | 7 | 3 |
| Disallowed (%) | 1 | 1 | 0 | 1 |
| Protein rotamer outliers (%) | 3 | 3 | 4 | 2 |
| Density map (contour 0.2) | 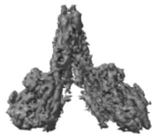 | 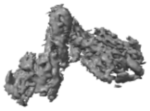 | 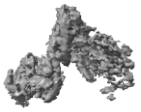 | 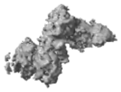 |
| Primary map (projection X) | 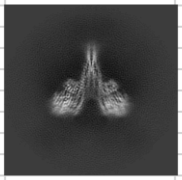 | 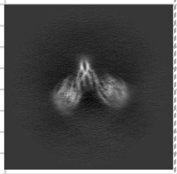 | 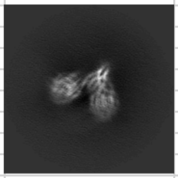 | 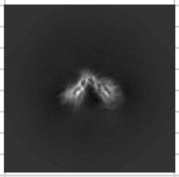 |
| Primary map (projection Y) | 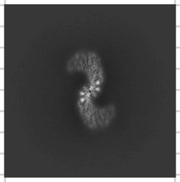 | 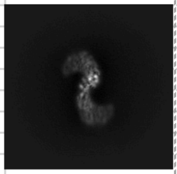 | 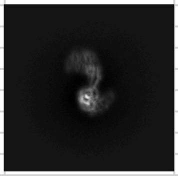 | 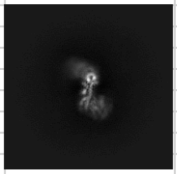 |
| Primary map (projection Z) | 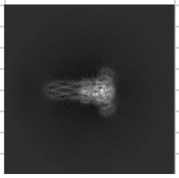 | 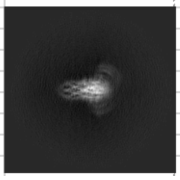 | 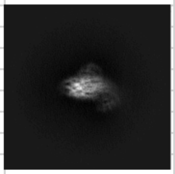 | 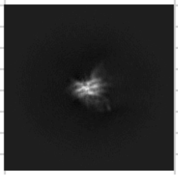 |

### Supporting Information:

Materials and Methods:

**IKK2 mutant and deletion constructs**

Full-length IKK2 (UniProt accession ID: O14920) with an N-terminal hexahistidine-tobacco etch virus (TEV) cleavage site tag was cloned in pFastBacHT B (Invitrogen) vector in frame within the BamHI and NotI sites. Other IKK2 gene constructs were subcloned similarly into pFastBacHT B for baculovirus expression.

IKK2 wild-type cDNA was inserted into pRCCMV-HA vector for transfection into mammalian cells to produce the HA-IKK2 fusion protein. Mutants of this vector were prepared by Q5 site-directed mutagenesis protocol (New England Biolabs).

The murine NF-κB p50 dimerization domain (DD), residues 245-358, was fused to HA-IKK2$^{FL}$, IKK2$^{1\text{-}669}$ and IKK2$^{MONO}$ (1-637 with an internal deletion 477-526) with a linker of 5XGlycine.

**Retroviral reconstitution of MEF cells**

HA-tagged wild-type and mutants of IKK2 were transferred from pRC-HA and cloned into the BamH1 and Not1 sites in the pBABE retroviral vector. HEK293T cells were transfected with 8μg of pBABE-IKK2 constructs in combination with 3μg of pCL-Eco plasmid in 6 cm petri dishes. Supernatant containing recombinant virus from these plates were collected 42 h post-transfection and filtered through a 0.45-micron filter. Previously seeded ikk2$^{-/-}$ MEF cells were infected with this virus at a ~50- 60% confluency along with 4 μg/ml of polybrene in 10cm plates., MEF cells were split into 15 cm plates after ~ 48 h of infection and selected with 3-5 μg/ml of Puromycin. When the selection was complete (about 3-5 days in presence of Puromycin), Puromycin resistant cells were selected after 3-5 days and compared for expression of IKK2 equivalent to wild-type MEF cells, and used for downstream signaling assays.

**Protein expression and purification**

His-tagged NEMO proteins were expressed in Rosetta (DE3) *E. coli* cells (MilliporeSigma). Cultures of 1 l LB media with 100 μg/ml ampicillin were grown at 37 °C to attenuance of 0.2 at 600 nm before induction with 0.2 mM IPTG, and stirred at 150 rpm for 16 h at 22 °C. Cells were harvested by centrifugation at 3000*g* for 10 min (Beckman Coulter), and cell pellets were lysed by sonication (VWR Scientific) on ice in 200 ml of lysis buffer (20 mM Tris–HCl [pH 8.0], 500 mM NaCl, 10% w/v glycerol, 10 mM imidazole, 0.2% Triton X-100, 1 mM PMSF, and 5 mM β-mercaptoethanol). Lysates were clarified by centrifugation at 30000g for 45 min. Supernatants containing soluble proteins were applied to a 1 ml Ni–NTA agarose column that was pre-equilibrated with the lysis buffer. Bound proteins were washed with 200 ml wash buffer (lysis buffer with 40 mM imidazole) and eluted in 10 ml elution buffer (lysis buffer containing 150 mM NaCl and 250 mM imidazole).

For purification of IKK2, respective Sf9 insect cells from 1 l suspension cultures were harvested by centrifugation at 3000*g* for 10 min at 4 °C and lysed by sonication in 100 ml of lysis buffer (25 mM Tris–HCl, pH 8.0, 200 mM NaCl, 10 mM imidazole, 10% w/v glycerol, and 5 mM β-mercaptoethanol). The lysate was clarified by centrifugation twice at 25,000g for 45 min at 4 °C. Pre-equilibrated Ni–NTA agarose resin was added at a ratio of 1 ml of resin slurry/liter of lysed cell culture, and the mixture was incubated on a rotator at 4 °C for 3 h. The Ni beads were pelleted at 600g for 2 min in a swinging bucket centrifuge rotor. Supernatant was carefully decanted, and the protein-bound resin was resuspended with wash buffer (lysis buffer containing 30 mM imidazole) and incubated at 4 °C on a rotator for 2 min. The Ni beads were pelleted again and decanted (wash 1). This was repeated until the last wash fraction contained 0.01 to 0.1 mg/ml of protein (Bio-Rad Protein Assay). Elution buffer (lysis buffer containing 250 mM imidazole) was added, and eluted fractions were collected and stored at −80 °C.

**Cryo-EM sample preparation and data collection**

For assembly of complexes, size-exclusion purified IKK2 (final ~2 μM) in 150 mM NaCl, 2 mM dithiothreitol (DTT), 50 mM Tris-Cl buffer, pH 7.5 and 4% glycerol was incubated with size-exclusion purified NEMO$^{1-110}$ (final ~ 4μM) at 4 °C for 15 min. The concentration of IKK2 and NEMO were determined by measuring absorbance at $A_{260}$ with a NanoDrop (Thermo Scientific). The concentration of the complex was then adjusted to ~1 μM with abovementioned buffer without glycerol so that final glycerol concentration is less than 2%. Complexes (4 μl) were then applied manually two to three times in quick succession to Quantifoil R 1.2/1.3 Au grids that had been freshly plasma cleaned in Gatan Solarus II (under Ar/$O_2$ mixture) for 10 s. After application, grids were blotted with varying force and times using a Vitrobot Mark IV (Thermo Fisher) at 4 °C and 100% humidity, and then immediately plunge-frozen in liquid ethane.

Cryo-EM data were collected on a 300-keV Titan Krios cryo-transmission electron microscope at Titan Krios cryo-transmission electron microscopes (FEI Company) at the University of California San Diego Cryo-EM facility equipped with GIF quantum energy filters attached to a Falcon 4 electron counting direct detector (Gatan) using EPU 2 (Thermo Fisher). Images were acquired at a magnification of approximately 130,000, corresponding to a calibrated raw pixel size of about 0.5 Å, in EFTEM mode with an energy filter slit width of 10 eV. Video image stacks with a total electron dose of roughly ~60 $e^{-}$/Å$^2$ were saved in non-super resolution counting mode.

**Cryo-EM data processing and analysis**

The collection and processing of raw data is outlined in detail in **Fig. S1 and S2**. Briefly, raw videos of each independent dataset were imported in cryoSPARC 3.1.1 (1), motion-corrected using patch motion correction (multi) of cryoSPARC and CTF estimated using patch CTF estimation (multi) in cryoSPARC. A list of micrographs for each dataset (removing poor quality micrographs using various criteria, for example, CTF fit of less than 9 Å, astigmatism less than 800, motion distribution less than 45 and motion curvature less than 30) was prepared. Data were collected at an ice thickness that did not lead to particle denaturation and produced minimal background noise. Particles were picked from curated micrographs using various methods and iterative strategies outlined in Extended Data Fig. 2. Duplicate particles were removed in cryoSPARC, and binned or un-binned particles were extracted from micrographs depending on the purpose and stage of analysis. Several rounds of two-dimensional classification were performed to eliminate poor quality particles that could not be classified (**Fig. S3**). Selected particles from two-dimensional classes were used for initial 3D classification (with different number of classes) using ab initio reconstruction. Particles belonging to poorly resolved ab initio 3D class averages were discarded. Particles from selected 3D classes were used for heterogeneous classification with various plausible models. Iteration of the above processes provided a refined set of particles, which were classified into probable 3D classes (judged from shape profiles of 3D models of heterogeneous classification), and eventually particles from final 3D class sets were extracted without binning with a 320-pixel box size for consecutive refinement using homogeneous, non-uniform (and local refinement if map improvement was observed) algorithms in cryoSPARC with appropriate masks. The total number of videos, number of particles used for final refinement of each class, orientation distribution, half-map-based correlation, global resolution estimate and several other parameters are provided in **Fig. S4**. Density sharpened versions of the maps were used for model building.

**Model building and refinement**

The model building pipeline is depicted in detail in **Fig. S5**. In brief, an initial model of IKK2 was built manually by further refining the deposited coordinate in the deposited electron density map (PDB 4KIK). The models (partial domains) were rigid body fitted models in the respective electron potential maps and iteratively (re)built in real space in Coot (v.0.9.8.1)(2). Since many regions are of high flexibility, the models were refined minimally in CCPEM (v.1.6.0)(3) to derived *B*-factors. empirically identified to provide the highest map correlation coefficient, as calculated with Phenix (v.1.18.2–3874)(3) (**Table S2**). Figures of molecular models were made with PyMOL (v.3.0.3, PyMOL Molecular Graphics System, Schrödinger, LLC) and ChimeraX (v.1.8)(4).
